# Sorted-Cell Proteomics Reveals an AT1-Associated Epithelial Cornification Phenotype and Suggests Endothelial Redox Imbalance in Human Bronchopulmonary Dysplasia

**DOI:** 10.1101/2025.03.20.644398

**Authors:** Mereena George Ushakumary, William B Chrisler, Gautam Bandyopadhyay, Heidie Huyck, Brittney L Gorman, Naina Beishembieva, Ariana Pitonza, Zhenli J Lai, Thomas L Fillmore, Isaac Kwame Attah, Andrew M. Dylag, Ravi Misra, James P. Carson, Joshua N Adkins, Gloria S. Pryhuber, Geremy Clair

**Affiliations:** Biological Sciences Division, Earth and Biological Sciences Directorate, Pacific Northwest National Laboratory, Richland, WA, 99352, USA; Environmental Molecular Sciences Laboratory, Earth and Biological Sciences Directorate, Pacific Northwest National Laboratory, Richland, WA, 99352, USA; Department of Pediatrics, University of Rochester School of Medicine and Dentistry, 601 Elmwood Avenue, Rochester, NY 14642, USA; Texas Advanced Computing Center, University of Texas at Austin, Austin, TX, USA

**Author notes:** Corresponding authors: Mereena George Ushakumary and Geremy Clair, Address: 3300 Stevens Dr, Biological Sciences Facility, MSIN J4-02 Richland, WA 99354.

**Keywords:** proteomics, epithelial cells, endothelial cells, BPD, lung, prematurity

## Abstract

Bronchopulmonary dysplasia (BPD) is a neonatal lung disease characterized by inflammation and scarring leading to long-term tissue damage. Previous whole tissue proteomics identified BPD-specific proteome changes and cell type shifts. Little is known about the proteome-level changes within specific cell populations in disease. Here, we sorted epithelial (EPI) and endothelial (ENDO) cell populations based on their differential surface markers from normal and BPD human lungs. Using a low-input compatible sample preparation method (MicroPOT), proteins were extracted and digested into peptides and subjected to Liquid Chromatography-tandem Mass Spectrometry (LC-MS/MS) proteome analysis. Of the 4,970 proteins detected, 293 were modulated in abundance or detection in the EPI population and 422 were modulated in ENDO cells. Modulation of proteins associated with actin-cytoskeletal function such as SCEL, LMO7, and TBA1B were observed in the BPD EPIs. Using confocal imaging and analysis, we validated the presence of aberrant multilayer-like structures comprising SCEL and LMO7, known to be associated with epidermal cornification, in the human BPD lung. This is the first report of accumulation of cornification-associated proteins in BPD. Their localization in the alveolar parenchyma, primarily associated with alveolar type 1 (AT1) cells, suggests a role in the BPD post-injury response. In the ENDOs, redox balance and mitochondrial function pathways were modulated. Alternative mRNA splicing and cell proliferative functions were elevated in both populations suggesting potential dysregulation of cell progenitor fate. This study characterized the proteome of epithelial and endothelial cells from the BPD lung for the first time, identifying population-specific changes in BPD pathogenesis.

**New & Noteworthy:** The study is the first to perform proteomics on sorted pulmonary epithelial and endothelial populations from BPD and age-matched control human donors. We identified an increase in cornification-associated proteins in BPD (e.g., SCEL and LMO7), and evidenced the presence of multilayered structures unique to BPD alveolar regions, associated with alveolar type 1 (AT1) cells. By changing the nature and/or biomechanical properties of the epithelium, these structures may alter the behavior of other alveolar cell types potentially contributing to the arrested alveolarization observed in BPD. Lastly, our data suggest the modulation of cell proliferation and redox homeostasis in BPD providing potential mechanisms for the reduced vascular growth associated with BPD.

## Introduction

Bronchopulmonary dysplasia (BPD) affects up to 50% of newborns born prematurely and is the most significant pulmonary morbidity of prematurity(1). Premature infants, less than 37 weeks gestation, are at risk for BPD inversely related to their gestational age and directly to the degree of pulmonary immaturity as they are born during early stages of lung development (late canicular or saccular) compared to term infants(2, 3). Neonatal exposures such as hyperoxia, mechanical ventilation, inflammation, and infection all contribute to lung injury, resulting in halted alveolarization, reduced vascular growth, and increased fibroblast proliferation in BPD patients(4). Despite advances in neonatal care, therapies for BPD remain inadequate and are mainly focused on providing support to the damaged lungs, allowing but not necessarily augmenting organ repair after neonatal lung injury. Therefore, it is important to understand the cellular and molecular pathogenesis of BPD to elucidate factors promoting its progression to target effective preventive and therapeutic strategies.

Proteomic studies have been conducted on various samples from premature infants, including plasma, urine, tracheal aspirates, and, more recently, pulmonary tissues(5–8). The first three sample types allow for the less invasive detection of disease-related proteins as potential diagnostics, yet they are peripherally associated with the disease and do not necessarily reflect the molecular mechanisms driving disease within the pulmonary tissues. Our group previously reported proteomics on human BPD lung tissues in different stages of disease in comparison to corrected age-matched control pediatric donors, quantifying 5,746 unique proteins and revealing cell-population shifts and tissue-level proteomic changes(8). This study, however, was limited by the complexity of proteomics on approximately 1 cm^3^ whole tissue blocks, leaving a gap in understanding the contribution of individual cell populations to the observed changes.

Here, we conducted the proteomic profiling of epithelial (EPI) and endothelial (ENDO) cell populations obtained through fluorescence-activated cell sorting (FACS) from dissociated lung tissue cell suspensions originating from both control and BPD human donors. The cell suspensions were prepared from freshly dissociated pulmonary tissue blocks prior to cryopreservation as part of the BioRepository for INvestigation of Diseases of Lung (BRINDL). BRINDL was created for the Developmental Lung Molecular Atlas Program (LungMAP) consortium and comprises samples from more than 470 pediatric and adult human cases(9). Similar BRINDL cell suspension preparations were previously used to perform flow sorted cell muti-omics analyses(10–12) and to generate primary cell culture models(13). Knowing the previously reported abnormalities of vascular and epithelial neighborhoods, growth and function that occur in BPD(14–17), we hypothesized that proteomics from enriched EPI and ENDO populations would reveal proteins associated with chronic tissue damage and repair in BPD, particularly those that drive cellular shape and stiffness. The novel and detailed data demonstrate the topmost proteins modulated in the BPD EPI population and include proteins known to contribute to cytoskeletal structures. More specifically, Sciellin (SCEL) and LIM domain only 7 (LMO7) were abnormally abundant in the BPD EPI sorted population. Both proteins were previously suggested to be involved in the cornification process in epithelia. Cornification is a specialized form of programmed cell death where epithelial cells undergo a process of terminal differentiation, losing their nucleus and organelles, and becoming filled with keratin proteins, resulting in a tough, protective, and essentially dead cell layer, most commonly seen in the outer layer of the skin (stratum corneum)(18). In the endothelial population, many of the BPD modulated proteins were related to redox homeostasis. These observations provide groundwork for further exploration in model systems to understand their significance in lung physiology and pathology.

## Materials and Methods

### Study population

The studied lung samples were authorized by next of kin for donation for research, when transplantation was not possible, through the United Network of Organ Sharing (UNOS) Organ Procurement Transplantation Network in coordination with the International Institute for Advancement of Medicine (IIAM) and National Disease Research Interchange (NDRI). The University of Rochester IRB approved and oversees the placement and management of these samples in BRINDL (RSRB00047606). The study reports on 10 human donor lungs: 5 healthy controls and 5 with clinical and histopathological BPD. The postnatal age of all donors was between 13 and 24 months corrected for gestational age at birth. Both groups included male and female donor tissues. The donor demographics are provided in the Supplementary Table 1.

### Lung cell dissociation

Lung cells were dissociated by mechanical disruption and enzymatic digestion as described previously(19–21). In brief, distal lung tissues from right upper and right middle lobes were scissor-cut into small pieces, the proximal tough airways removed, and remainder placed in c-tubes (Miltenyi Biotech) with 10 ml/tube digestive enzyme cocktail containing 2 mg/ml collagenase A (Roche), 1 mg/ml dispase II (Gibco), 0.5 mg/ml elastase (Worthington Biochemical), and 2 mg/ml bovine pancreas deoxyribonuclease-I (DNase-I, Sigma). Tissues were mechanically disrupted by GentleMACS^®^ Octo Tissue Dissociator (Miltenyi Biotech) and then incubated with digestion cocktail in a 37 °C incubator for an hour with occasional mixing. After incubation, the enzymatic reaction was stopped by addition of ice-cold neutralization buffer (5 ml/tube with 10% FBS [Atlanta Biologicals] containing Dulbecco’s Phosphate Buffered Saline [Lonza BioWhittaker]). Digested tissues were then passed through 100-micron strainer (Falcon, Corning), centrifuged, buffer containing digestive enzymes removed, and the cell pellet treated with 20 ml/tube ACK buffer (Lonza BioWhittaker) for 5 minutes at room temperature to remove any contaminating red blood cells. ACK buffer was neutralized by addition of up to 50 ml cold sterile DPBS containing 10% FBS followed by centrifugation. The pelleted cells were then resuspended, counted by trypan blue exclusion, washed in cold DPBS and resuspended in freezing medium, (10% DMSO (Sigma) and 90% FBS, ∼50×10^6^ cells/ml), slow cooled and stored in liquid nitrogen until further processing.

### Fluorescent activated cell sorting

Fluorescence activated cell sorting was performed as described earlier(19, 20). In brief, frozen mixed-lung cells were thawed, washed, pelleted, and resuspended in 1 ml 1x PBS. Postfreeze viable cell counts were determined by hemocytometer and trypan blue exclusion. Cells were then incubated in 1x PBS with FcR blocking reagent (Miltenyi Biotech, Cat # 130-059-901) to block nonspecific Fc-receptor binding. Cells were washed after Fc-receptor blocking and incubated at a final concentration of 1L×L10^6^ cells per 10 μl of staining-specific antibodies: CD326/EpCAM monoclonal antibody (eBioscience, Cat # 53-9326-42, Lot # 2563945), CD31/PECAM antibody (Miltenyi Biotech, Cat # 130-128-241, Lot # 1323071017) and CD144/VE-Cadherin antibody (Miltenyi Biotech, Cat # 130-125-985, Lot # 1323071125) for 90 min at 4 °C, protected from light. The cells were then washed x 3 and resuspended in PBS before passage through a 70-μm strainer (Corning). Final mixed cells were suspended in 1 ml of 1X PBS before sorting.

### Protein extraction and digestion

Protein extraction and digestion was performed by the microscale proteomic method called microdroplet processing in one pot for trace samples (microPOTS)(22). Briefly, cell pellets were treated with 0.05% n-dodecyl-b-D-maltoside (DDM) and dithiothreitol (DTT) and incubated at 37 °C for 30 minutes followed by 10 mM Iodoacetamide (IAA). Digestion was done with 0.01 μg/μL trypsin and incubated overnight. Formic acid was then added at 1% final concentration and the sample was diluted to 35 μL with water. Protein from each sample (25 μL of 0.01 μg/uL) was submitted for LC-MS/MS analysis.

### Liquid chromatography-tandem mass spectrometry

MS analysis was performed using a Q-Exactive HF-X mass spectrometer (Thermo Scientific) by acquiring datasets for liquid chromatographic separation of a 0.1 ug/uL of peptide solution. The ion transfer tube temperature and nano electrospray voltage were 300 °C and 2.2 kV, respectively. Each dataset was collected for 120 min following a 10 min delay after completion of sample trapping and start of gradient. FT-MS spectra were acquired from 300 to 1800Lm/z at a resolution of 60Lk (AGC target 3e6) and the top 12 FT-HCD-MS/MS spectra were acquired in data-dependent acquisition (DDA) mode with an isolation window of 0.7 m/z at a resolution of 45 k (AGC target 1e5). A normalized collision energy of 30 was used for HCD fragmentation, with a 45 s exclusion time, analyzing only charge states 2 to 6. Liquid chromatography separations were done using a Thermo Dionex Ultimate, configured with 2 pumps for sample trapping and reverse-flow elution of the sample onto the analytical column respectively.

### Proteomics data analysis and statistics

Proteomics raw data files were analyzed using MsFragger (v20.0)(23) using the LFQ-MBR workflow with the default settings against the UniProt non redundant human proteome database (downloaded in December 2023, 20,654 sequences). The resulting quantitative data was further processed using the package RomicsProcessor v1.1 (https://github.com/PNNL-Comp-Mass-Spec/RomicsProcessor). The data was log2-transformed and median normalized. Data were not imputed to avoid the introduction of systematic quantitative biases, two types of statistics were performed: Student’s T-test was used to compare the protein abundance between groups (when at least 50% of values were not missing for either group and the data was normally distributed), binomial generalized linear model (GLM) tests were used to identify binary protein presence or absence between groups. Enrichment analyses were performed using the tool Protein MiniOn v0.3.0 (https://github.com/GeremyClair/Protein_MiniOn) using Gene Ontologies (GO)(24), KEGG(25) and Reactome pathway(26) downloaded from the databases alongside the Fasta file (in December 2023) using the EASE score, a modified Fisher’s exact test developed for DAVID (8). The Massive ID to access the raw dataset is MSV000094373.

### Immunofluorescence staining and analysis

Immunofluorescent staining was performed on 5-µm-thick sections from paraffin embedded lung tissue blocks from the same donors represented in the dissociated cell studies. Slides were deparaffinized in xylene, rehydrated in a series of graded ethanol and washed in 1x PBS. When required, antigen retrieval in 10 mM citrate buffer (pH 6.0, Abcam) was performed. Non-specific antibody binding was blocked with 4% normal donkey serum (Abcam) in PBS with 0.1% Triton X-100 (PBST) for 2 hours. Slides were incubated in primary antibodies (Table 2) diluted in blocking buffer overnight at 4 °C. After washing in PBST, slides were incubated in fluorescent secondary antibodies (1:200, Invitrogen) and DAPI (1 µg/ml, ThermoFischer Scientific) diluted in blocking buffer, for 1 hour at room temperature. Slides were subsequently washed in PBST and mounted in Prolong Gold (ThermoFisher Scientific). Images were captured on a Leica Epifluorescence microscope.

**Table 1:** Donor Demographics.

| Donor Id | BPD or CNT | GA at Birth (weeks) | Corrected Age (months) | Sex | Race | Cause of Death |
| --- | --- | --- | --- | --- | --- | --- |
| D119 | CNT | 40 | 13.94 | M | White | Meningitis with severe brain injury |
| D090 | CNT | 40 | 14.46 | F | Unknown | Gunshot wound |
| D079 | CNT | 40 | 15 | M | White | Blunt head trauma |
| D006 | CNT | 40 | 19.42 | F | White | Drug intoxication |
| D095 | CNT | 40 | 21.58 | F | White | Head trauma |
| D184 | BPD | 24 | 11.47 | M | White | Cardiac arrest |
| D227 | BPD | 23 | 16 | F | White | Anoxic brain injury |
| D115 | BPD | 26 | 16.13 | F | White | BPD with pulmonary hypertension |
| D170 | BPD | 25 | 18.3 | F | Black | BPD with clinical severe pulmonary hypertension |
| D401 | BPD | 26 | 34.23 | M | White Hispanic | Chronic lung disease |

BPD, Bronchopulmonary dysplasia; Corrected Age, post-natal age + (gestational age – 40 weeks); CNT, Control; GA, Gestational age; F, Female; M, Male.

**Table 2.**
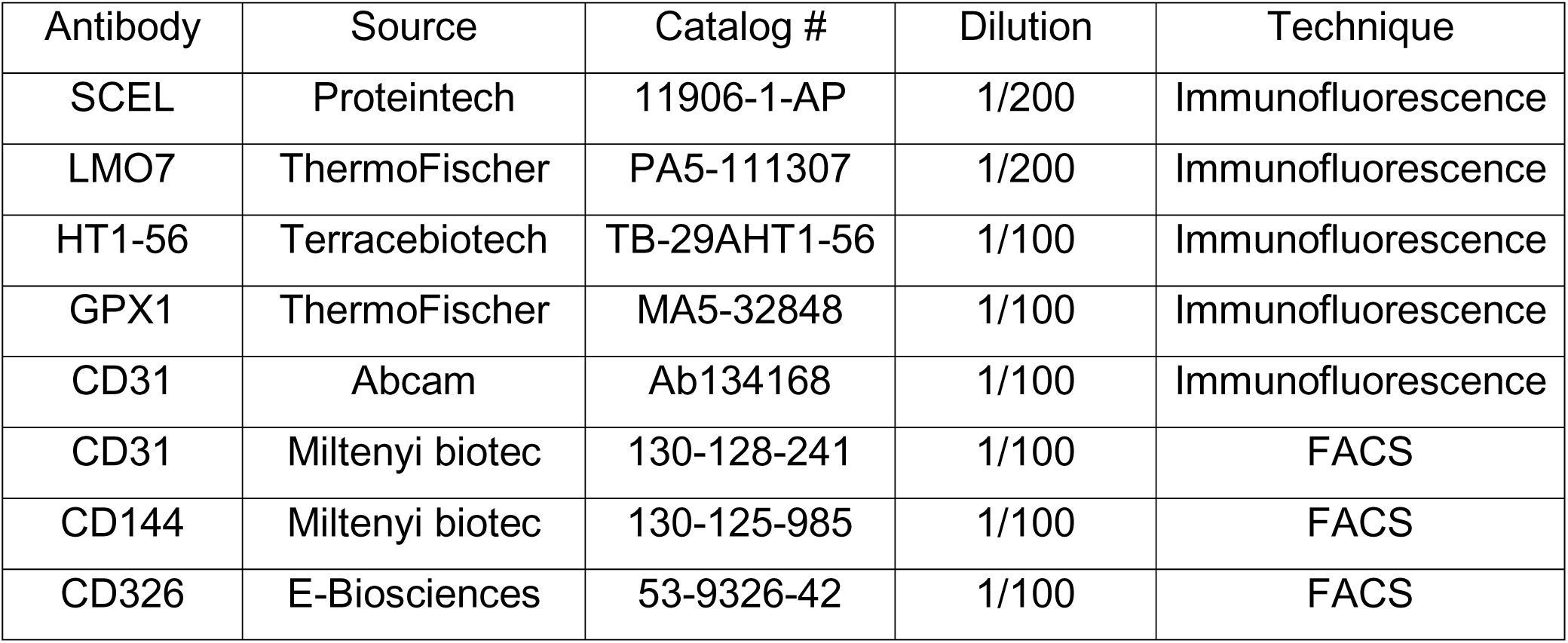
List of the primary antibodies used in this study.

Additional images were acquired on a Zeiss LSM 710 Confocal microscope using laser excitation wavelength 405 nm, emission 410-535 nm for DAPI; excitation 488 nm, emission 490- 557 nm for LMO7 and SCEL; and excitation 561 nm, emission 570-660 nm for HT1. The EC Plan-Neofluar 20x/0.50 objective was used to capture 10 fields of view (FOV) per tissue that were further analyzed using FIJI - Image J software. The quantification was done based on a previously published protocol.(27) The ratio of SCEL to DAPI (SCEL/DAPI) or LMO7 to DAPI area (LMO7/DAPI) was calculated and averaged across 10 FOV per donor. The ratio values were compared between healthy and diseased samples using a two-tailed T-test.

## Results

### FACS sorting of epithelial and endothelial cells from BPD Lung dissociated cells

Dissociated cells were enriched based on their differential surface protein markers. The dissociation, sorting and staining strategies used are outlined in Fig.1. Epithelial cells (EPIs) were selected by the presence of EpCAM at their surface. Endothelial cells (ENDOs) were selected based on the presence of CD31 and CD144 at their surface. The number of EPIs and ENDOs was evaluated based on the percentage of parent dissociated cell population (% parent). The % parent of the BPD EPIs was not significantly different from controls (Fig.2a). However, the % parent of the ENDOs obtained from the BPD samples was significantly reduced compared to the controls (Fig.2b). These results, in line with the existing literature, suggest reduced vascular development or the reduction of the vascular bed in the BPD lungs(8, 28, 29).

**Figure 1.**
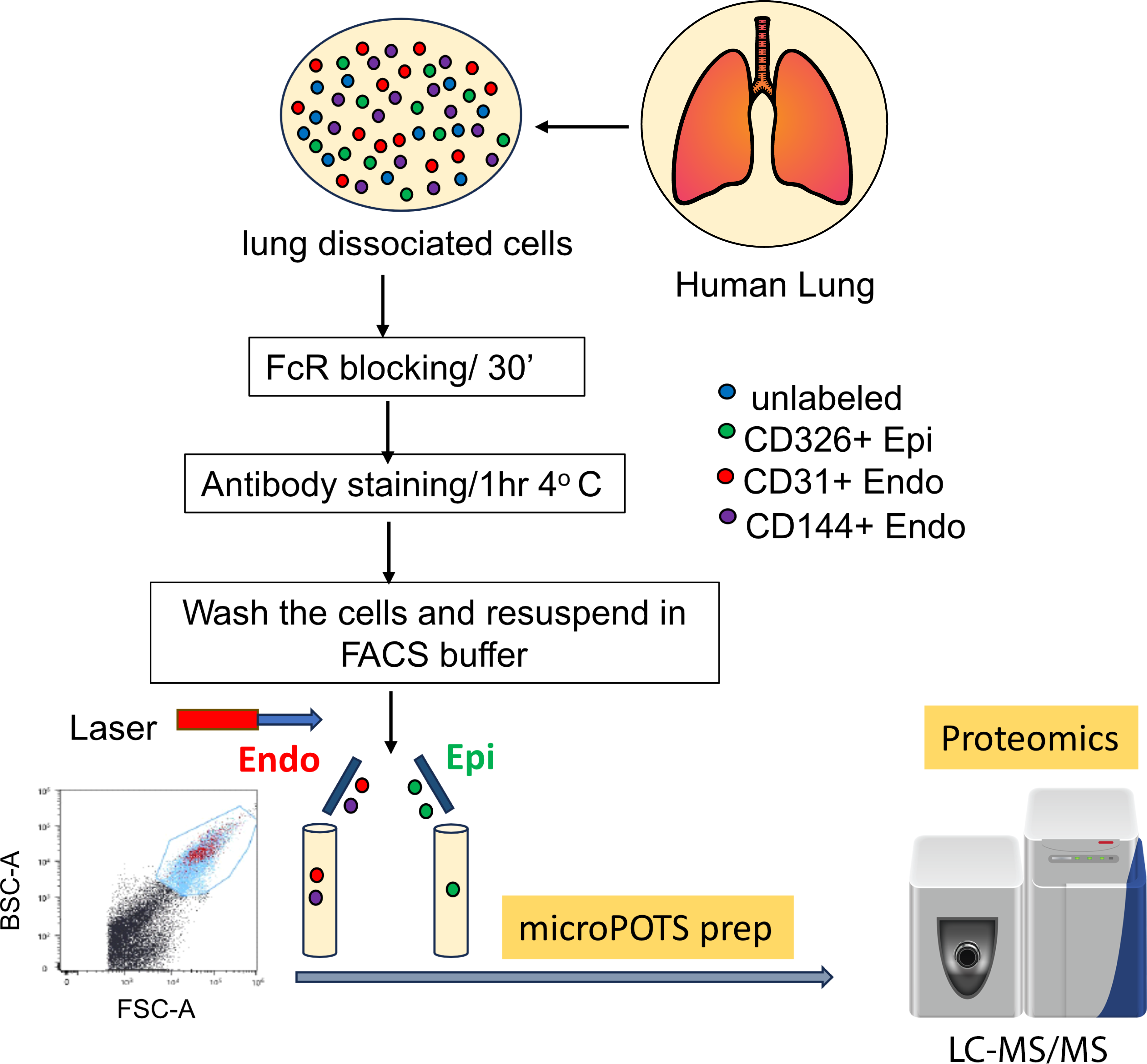
Schematic representation of human lung tissue dissociation, cell sorting strategy and proteomics. The dissociated cells from 10 corrected age-matched donors (n=5 full-term born controls, n=5 BPD) were sorted for EpCAM+ (CD326+) epithelial cells and CD31+CD144+ endothelial cells. Sorted populations were prepared for LC_MS/MS proteomics analysis using microPOT sample preparation protocol.

**Figure 2.**
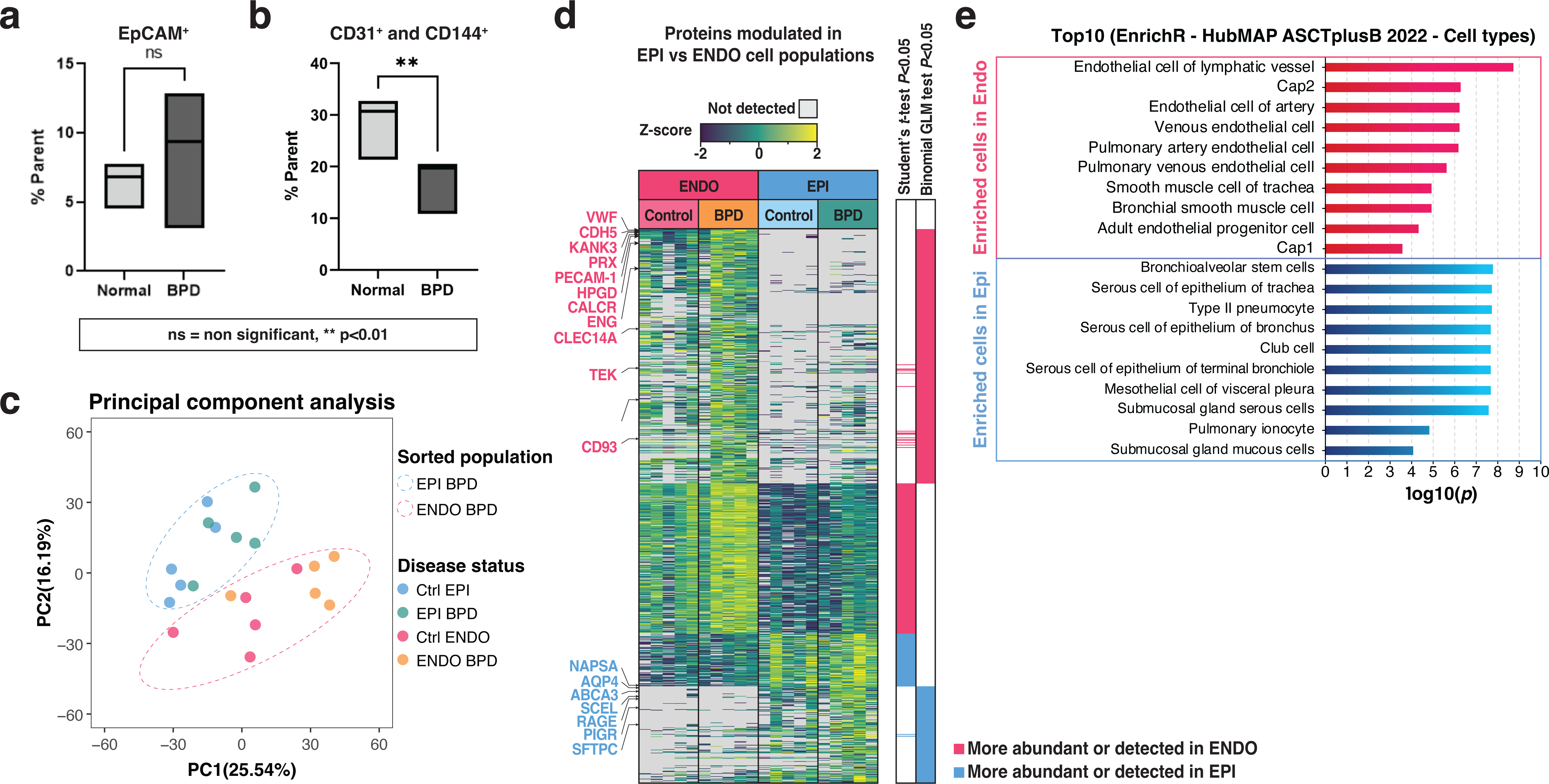
Analysis of sorted epithelial and endothelial cell populations from control and BPD lungs. (a) The percentage of sorted EpCAM+ cells in the parent unsorted population was calculated. The graph shows no significant difference in sorted BPD epithelial population from control lungs; ns- not significant. (b) The percentage of sorted CD31+CD144+ endothelial populations were significantly decreased in BPD lungs compared to control lungs; \*\**p*<0.005 by Student’s t-test. (c) Principal component analysis (PCA) plot showing the separation between samples by color for cell type and disease status. (d) Heatmap depicting the proteins modulated in abundance or detection in the BPD and control lungs for both epithelial and endothelial sorted populations. The statistics bar indicates p< 0.05 by Student’s T-test and Binomial GLM test used to evaluate abundance changes as well as presence/absence, respectively. (e) Results of enrichment analysis performed using EnrichR against the HubMAP ASCT+B database, confirming the identity of the sorted cell populations.

### Proteome profiling confirms purity of sorted EPI and ENDO cell populations

A total of 4,970 proteins were detected, of these 2,043, were detected in at least 50% of the EPI or ENDO sorted population samples (regardless of disease status) and were considered quantifiable. The principal component analysis (PCA) plot shows the segregation of proteome profiles of the EPI and ENDO sorted populations (Fig.2c). The proteomic data were analyzed using Student’s T-tests to estimate abundance difference between groups. Binomial GLM tests were used to evaluate binary differences in protein detection (protein presence/absence in the different groups), the postulate is that a protein with significantly higher detection (missingness not at random) is either more abundant or uniquely present in one of the two groups being compared. Scaled abundance (Z-scores) for the proteins in the EPI and ENDO groups regardless of disease status (test *p*<0.05) are depicted on the heatmap Fig.2d (quantitative data are provided as Supplementary Dataset 2). We found 246 proteins either significantly more abundant or detected in the EPI populations, 670 were more abundant or detected in the ENDOs (Supplementary Dataset 3). We used the EnrichR(30) (accessed December 2023) against the HuBMAP respiratory system ASCT+B augmented tables (2022)(31) to identify the pulmonary cell types enriched in each population based on cell-type markers (Fig.2e, Supplementary Dataset 4). As anticipated, data from EPI population were found enriched in pulmonary epithelial cells including multiple alveolar and airway cell types (Fig.2-e). For instance, AGER/RAGE and AQP4, known markers of AT1 cells as well as SFTPC, ABCA3, and NAPSA, known markers of AT2 cells, were enriched in the EPIs (Fig.2d). Multiple airway cell type markers such as PIGR, AGR2, or KRT5, were also enriched in the Epi population(32–34). All have been previously identified to be enriched in epithelial cells at the transcriptomics level (32, 33, 35). Similarly, the ENDOs were enriched in proteins known to be present in pulmonary lymphatic cells, capillary type 1 (Cap1) and 2 (Cap2) cells, as well as arterial and venous endothelial cell markers. The enrichment suggested that some smooth muscle cells may have been captured in the ENDO population likely due to their tight association with the endothelium. Proteins enriched in the ENDOs included PECAM1, VWF, CD93, TEK, CLEC14A, CLDN5, ENG, PRX, HPGD, CALCRL, CDH5, or KANK3 (Fig.2d). Overall, this data indicates that the sorting process worked as intended to obtain relatively pure epithelial and endothelial populations and confirms that protein expression profiles match reports of RNA expression from scSeq datasets(33–35).

### Proteome profiling of EPI and ENDO populations indicate disease related, cell population-specific, alterations in protein abundances

Between the EPIs originating from BPD donors and their aged-matched controls counterpart, 1,522 proteins were quantifiable (Supplementary Dataset 5), of which 293 were significantly modulated (either in detection or in abundance): 186 proteins were more detected (Binomial GLM *p*<0.05) in the BPD EPIs, 16 less detected, 88 more abundant (Student’s T-test *p*<0.05), and three less abundant (Supplementary Dataset 6).

In the ENDO population, of the 1,825 quantifiable proteins (Supplementary Dataset 7), 422 were significantly modulated either in abundance or in detection, 290 were more detected, 13 were less detected, 133 were more abundant, and 5 were less abundant (Supplementary Dataset 8).

### Epithelial-cell specific alterations suggest functional and structural changes in the BPD alveolar epithelium

The annotated heatmap shown in Fig.3a depicts the proteins differentially abundant or detected between the BPD and control EPI sorted populations; quantitative data are provided in Supplementary Dataset 5. Pathway enrichment analysis identified that the proteins more abundant in the BPD EPI population were related to cell proliferation and RNA splicing (Fig.3b, the full list is provided in Supplementary Dataset 9). Conversely, proteins lower in abundance/detection were enriched for terms such as “complement activation” (GO_BP), and “nitrogen metabolism” (KEGG). The GLM binomial test indicated that proteins more detected in BPD included: GOSR1 (Golgi-SNAP receptor complex 1), a membrane trafficking protein which transports cargoes between the endoplasmic reticulum and the Golgi and between Golgi compartments(36), supervillin (SVIL) a protein involved in mitosis reported to promote epithelial to mesenchymal transition(37), and prothrombin (F2), involved in blood coagulation, inflammation and wound healing. Two tight junction proteins, occludin (OCLN) and tight junction protein 2 (TJP2/ZO2), were more detected in the BPD compared to control EPIs. In addition, many proteins modulated in abundance or detection were associated with actin cytoskeletal functions, including TBA1B, DSG2, SCEL, and LMO7. Tubulin 1B (TBA1B) is known to play a critical role in the formation and stability of microtubules involved in maintaining cell shape, enabling intracellular transport, and facilitating cell division(38, 39). Desmoglein 2, DSG2, is a protein involved in skin cornification and shown to confer a hyperproliferative and apoptosis-resistant phenotype to skin epithelial cells(40). The role of SCEL and LMO7 will be discussed in the next section. Finally, less abundant/detected proteins in the BPD EPI population include mitochondrial proteins such as UQCRFS1, GSR, or SLIRP suggesting an altered energy metabolism in these cells.

**Figure 3.**
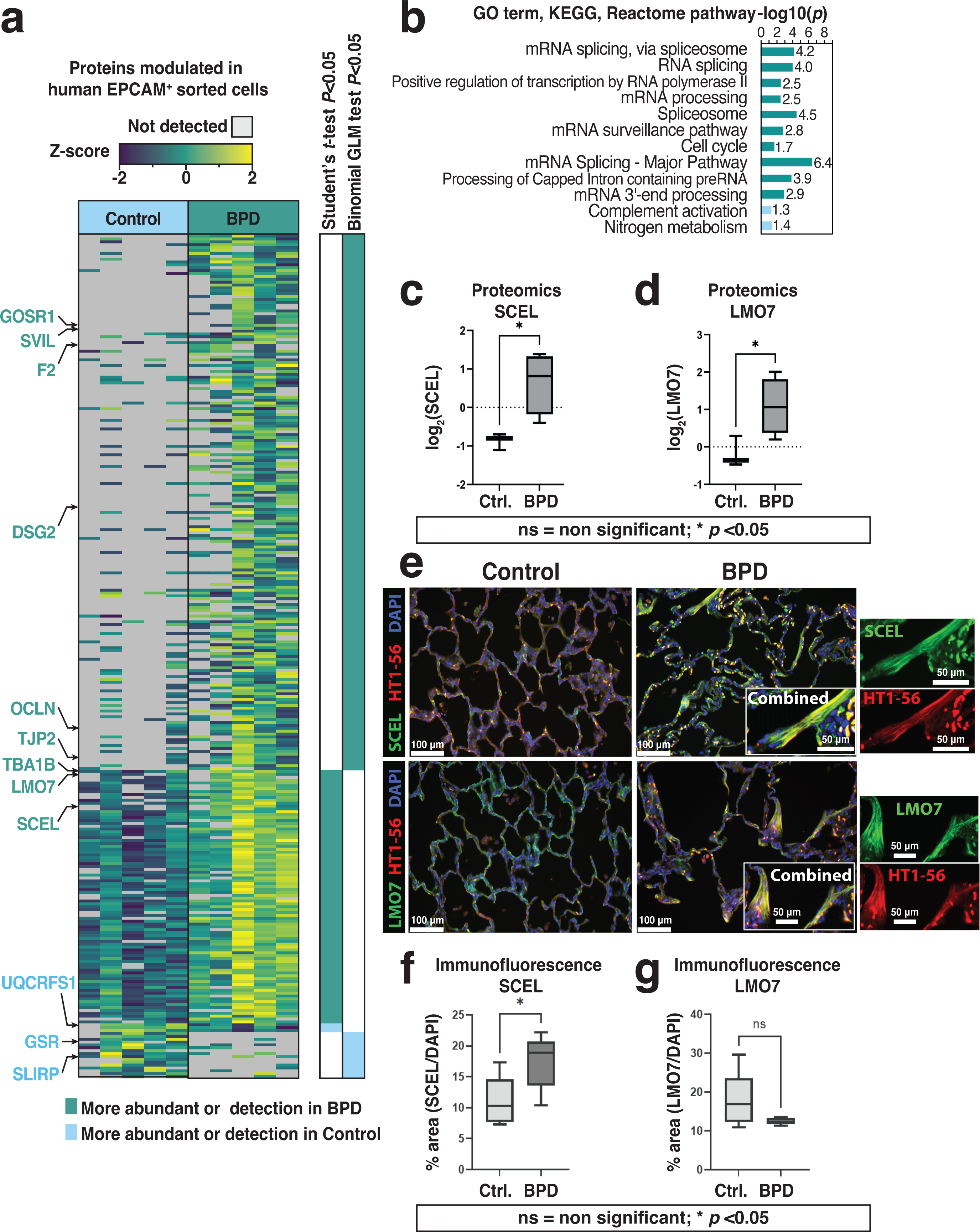
Epithelial cell protein dynamics in human BPD. (a) Heatmap of disease-modulated proteins in EPCAM+ cells. The statistics bar indicates p< 0.05 by Student’s T-test and Binomial GLM test used to evaluate abundance changes as well as presence/absence, respectively (b) Enrichment analysis of modulated proteins showing Gene Ontology (GO) terms cellular compartment (CC) as well as KEGG and Reactome pathways. Student’s t-test and binomial GLM test were performed to analyze more abundant/detected proteins in BPD epithelial cells compared to control. Enrichment of proteins with more abundant/detected and less abundant/detected are shown. (c) & (d) The protein abundance of the cornification-associated cytoskeletal proteins, SCEL and LMO7, represented by log_2_(LFQ); \**p*< 0.05. (e) Epifluorescence images of control and BPD donor lungs (FFPE, 5 μm, 20x magnification) showing SCEL and LMO7 staining with AT1 cell membrane marker HT1-56 and DAPI stained nuclei. (f) Quantification of SCEL and LMO7 immunofluorescence was performed by measuring the area covered by SCEL or LMO7 divided by the area of the nuclei (DAPI). ns= not significant, *p <0.05.

### The formation of a multilayer-like epithelial structure in BPD, comprising SCEL and LMO7, is associated with AT1 cells

As described above, SCEL and LMO7 were among the most significantly modulated protein in the BPD EPI population (Fig.3c-d) compared to the control EPI. SCEL is known to be involved in the formation of cornified envelopes(41). LMO7 is a microfilament-associated transcription factor. In the human Lung CellRef single-cell RNA sequencing dataset(33) SCEL and LMO7 are both found elevated in the AT1 cell cluster compared to other cell types. The CellRef UMAPs depicting their expression are provided as Supplementary Fig.1a-c & their distribution boxplots are shown in Supplementary Fig.1d&e (https://research.cchmc.org/pbge/lunggens/LGEA_v3_home.html, consulted in December 2024). Previously, using proteomics on whole tissue blocs, we found a significant reduction in the abundance of AGER, another AT1 marker and using multiplexed immunofluorescence, we have evidenced the increase of the proportion of SFTPC^+^ AT2 cells in the alveolar parenchyma by measuring the ratio of SFTPC^+^/panCytokeratin^+^ cells(8). While our previous results suggested a reduction of the AT1 population in BPD, we were surprised to observe an increase in SCEL and LMO7 abundance. We hypothesized that these proteins could contribute to the formation of cornified envelopes in the alveolar space in response to the BPD injury. To verify this hypothesis, we performed immunofluorescence (IF) imaging of SCEL and LMO7 along with HT1-56, an antibody previously described to stain exclusively AT1 cells in the human alveolar parenchyma(42, 43) (Fig.3e). Both SCEL and LMO7 were found in distribution consistent with AT1 in the control samples. In BPD exclusively, both proteins were sporadically observed to be abundant in multilayer-like structures of the alveolar epithelium, consistent with the architecture of a cornified epithelium. In addition, IF imaging demonstrates HT1-56 localized to the surface of these multilayered structures, suggesting that AT1 cells expressing the HT1-56 target protein contribute to the formation of these cornified-structures. Quantification of SCEL as the area of the protein divided by the area of the nuclei (DAPI) supports the proteomics result indicating a higher abundance of SCEL in the BPD epithelium (Student’s T-test, *p*<0.05, Fig.3f). However, quantification of LMO7 by immunofluorescence did not show significance between disease and control. The area covered by this protein was detected even in control tissues, in areas consistent primarily with AT1 localization. The additional presence of LMO7 in the squamous structures was not sufficient to reach significance (Student’s T-test, *ns*, Fig.3e & g) potentially being offset by the AT1 population being reduced in BPD and by the gene being expressed in other non-AT1 populations (see Supplementary Figure 1c).

### Proteomic landscape of the endothelial cells suggests redox imbalance

Our proteomic analysis revealed several significantly altered proteins in the BPD ENDOs compared to the aged match controls. (Fig.4a, Supplementary Dataset 7). Terms and pathways enriched in the proteins more abundant or more detected in the ENDO BPD population comprised “mRNA processing and splicing”, “positive regulation of translation” and “iron-sulfur cluster assembly” (Fig.4b, Supplementary Dataset 9). Terms and pathways enriched in the proteins less abundant were “innate immune response”, “collagen catabolic process”, “neutrophil degranulation” and multiple terms related to superoxide generation. We also observed changes in endothelial markers suggesting a potential shift in the cell types composing the ENDOs population in BPD. For example, the complement component C1q receptor (CD93/C1QR1) and the angiopoietin receptor (TEK/TIE2), two cell surface markers known to be expressed in some of the endothelial cell types but with low expression in Cap2 cells(33) (Supplementary Fig.2a-c), were found more detected in BPD. The gene expression distribution boxplots were also shown in Supplementary Fig.2d&e. Likewise, CDH5, ESAM and CLEC14A (Supplementary Fig.3a-c) were significantly more abundant or detected in BPD suggesting changes either in the proportion of endothelial cell types or altered endothelial cell states reflected in shifts in protein content. The expression distribution data in boxplot form is also displayed in Supplementary Fig.3d & e . In addition to endothelial specific proteins, many proteins involved in redox processes were altered in abundance or detection in the BPD ENDOs. Cytochrome b-245 light chain (CYBA), expected to increase ROS production, and superoxide dismutase (SOD1), expected to dismutase superoxides, were significantly less detected in BPD. On the other hand, many proteins involved in response to oxidative stress were more abundant or detected including: Nostrin, GSTK1, GLRX5, PRDX1, IFI30, the thioredoxin domain containing proteins TXNDC5 and TXNDC12, and GPX1. Notably, multiple cells in addition to CD31 positive endothelial cells were identified by immunofluorescence to be GPX1 positive preventing cell type specific quantitation (Fig.4c).

**Figure 4.**
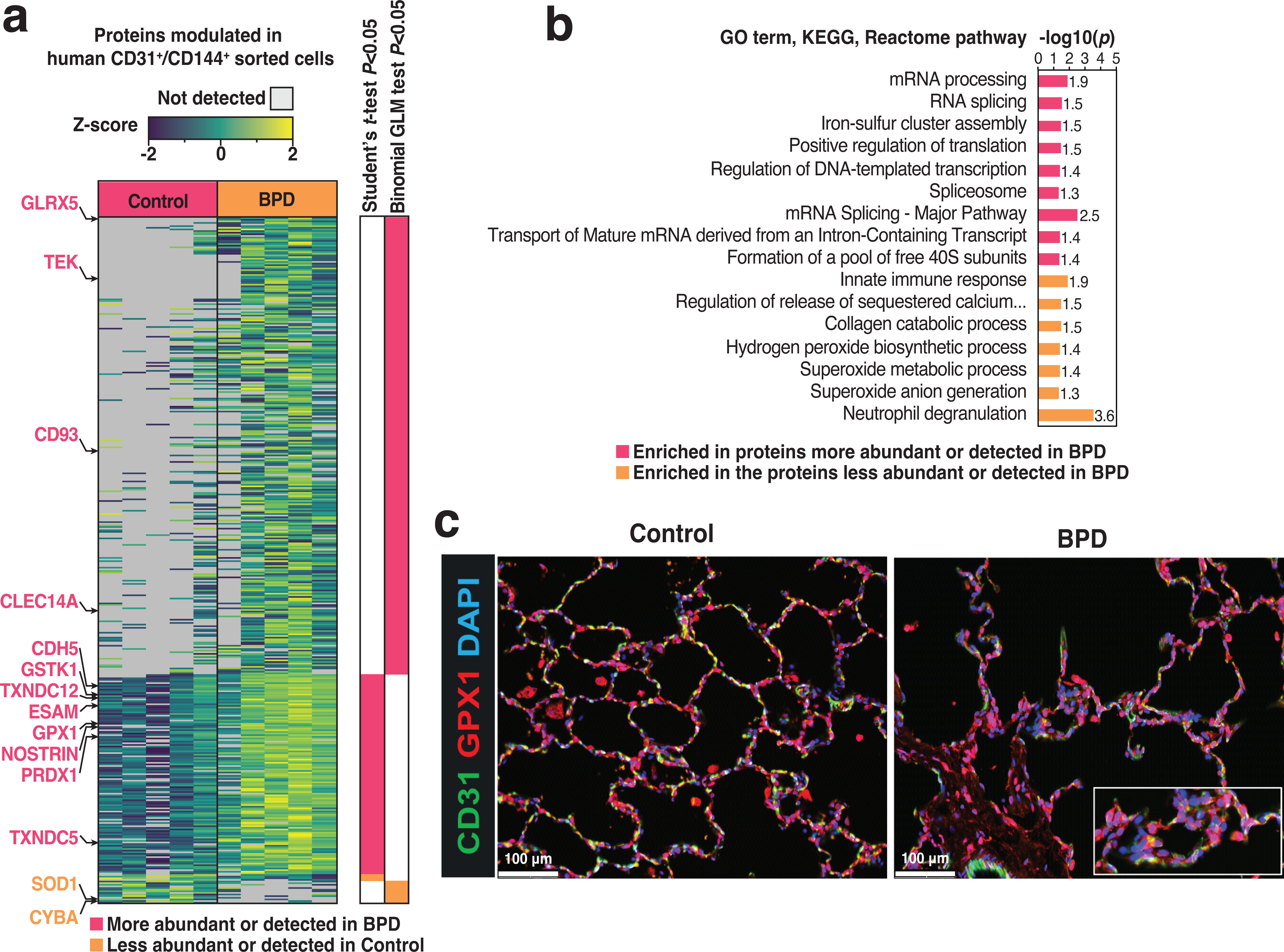
Endothelial cell protein dynamics in human BPD. (a) Heatmap showing the modulated proteins in CD31+ CD144+ endothelial cells. The statistics bar indicates p< 0.05 by Student’s t-test and binomial GLM tests. (b) Enrichment analysis of modulated proteins showing GO terms CC, BP (Biological process) as well as KEGG and Reactome pathways. Student’s t- test and binomial GLM test were performed to analyze more abundant/detected proteins in BPD endothelial cells compared to control. Enrichment of proteins with more abundant/detected and less abundant/detected are shown. (c) Epifluorescence images of control and BPD donor lungs (FFPE, 5 um, 20x magnification) showing staining for GPX1 with endothelial marker CD31 and DAPI stained nuclei.

## Discussion

While the roles of specific cell types in BPD has been explored using in vitro and animal models(44–52), the data generated from humans is limited due to the scarce availability of human BPD tissues. Our collaboration previously reported bulk proteomics alongside multiplexed immunofluorescence from normal and BPD donors(8). Recently, we published the proteomics and transcriptomics profiling of the four main cell lineages composing the lung to explore the cellular and molecular changes occurring during normal pulmonary post-natal development(12). We were able to identify protein-level markers of specific cell-types and to identify pathways that were specifically modulated in each cell lineage(12, 53). Similar studies, describing cell-lineage-specific changes in the BPD lung are lacking. We are now filling this gap by applying LC-MS/MS-based proteomics to characterize the proteome of sorted epithelial and endothelial cells populations from BPD-affected human donors and age-matched controls and highlight several novel observations.

The proteomics data generated evidenced that both the epithelial cells and endothelial cells had higher expression of the spliceosome components. Alternative splicing is generally elevated within stem cells and has been noted to contribute to pluripotency maintenance and differentiation(54). In addition, many mitosis-related proteins were found elevated in BPD suggesting further that the proportion of cells in a stem-like/plastic state might be higher in these cell populations in BPD. In the epithelial compartment, a population of bipotent AT2 is known to give rise to both AT1 and AT2(55). We have previously noted the increase of the AT2/AT1 proportion based on the increase of the SFTPC expressing epithelial cells(8), at least in part due to the reduced capacity of the AT2 bipotent progenitors to transition to AT1s(55, 56). Surprisingly, two of the proteins increased in abundance in our proteomics of BPD were SCEL and LMO7. Both SCEL and LMO7 are actin-associated proteins known to be involved in the formation of a cornified envelope in the skin epidermis. Cornification is a specialized type of programmed cell death where the epithelial cells lose their nucleus and organelles, and become filled with keratin protein, resulting in a tough, protective, and essentially dead cell layer, most seen in the outer layer of the skin(57). SCEL was previously identified as a marker of AT1 cells(58) and, in normal tissues, LMO7 expression is also enriched in AT1. In BPD, LMO7 expression is more confined to the regions with abnormal alveolar morphology. While the overall LMO7 protein level was elevated, reflecting higher abundance in these abnormal cells as indicated by the proteomics data, the increase was not found significant by imaging. This protein has been shown associated with cell junctional complexes and to shuttle between the plasma membrane and the nucleus in muscle cells(59). In epidermal cells, LMO7 was found to be strongly upregulated during cell differentiation to cornified keratinocytes(60) and was elevated during oral wound healing(61). Interestingly, it is a known TGF-β target gene yet a negative feedback regulator of TGF-β signaling and fibrosis(62). In addition to LMO7 and SCEL, we observed an increase in abundance of two tight-junction proteins: TJP2 and OCLN. Of note, tight-junctions proteins are known to remain in the cornified envelope during the cornification process(63). Finally, DSG2, another protein involved in cornification was found elevated in BPD. Exploring the hypothesis that cornified-like structures could be induced in the lung in response to the mechanical or oxidative stress sustained perinatally, using IF, we found existence of multilayered structures comprising LMO7 and SCEL tightly associated with AT1 cells based on their squamous shape and the expression of the antigen of HT1-56. Nuclei are not readily found in these structures consistent with a cornification phenotype. Finally, we observed the presence of Keratin 14, a type I intermediate filament found in healthy airway basal cells and identified in alveolar epithelial injury/disease(64) in regions enriched with SCEL (Supplementary figure 4), supporting a cornification response to injury. Cornification can be expected to contribute to reduced compliance and increased susceptibility to infection and inflammation. The role of our described AT1 cell phenotype in the pathophysiology of BPD, especially to arrested alveolarization and fibrogenesis remains to be explored.

While a reduction in endothelial populations is well demonstrated in BPD, the changes in cell-type proportions and function composing this “altered” endothelium remain to be elucidated. A recent transcript-level study performed scRNAseq and spatial transcriptomics from former preterm human infants with evolving BPD, and infants with both BPD and pulmonary hypertension (PH) evidenced the existence of unique aberrant capillary cell-states primarily in infants with BPD and PH(65). Here, using proteomics, we observed the increased ENDO abundance of CD93 and TEK, two cell surface markers expressed in some endothelial cell types reported to be low in the Cap2s (also known as “aerocyte” capillary cells). This result suggests changes in the Cap1/Cap2 ratio or could indicate the presence of cells in an aberrant capillary cell-state as suggested. We report many more endothelial specific markers to be modulated suggesting further studies should explore relationships to disturbances observed in BPD pulmonary endothelium.

Many redox proteins were identified to be modulated in the ENDO BPD samples. Nostrin, a regulator of the activity of the endothelial nitric oxide synthase (eNOS)(66), GSTK1 which is known to conjugate glutathione to exogenous and endogenous compounds, GLRX5, a protein involved in the biogenesis of iron-sulfur clusters, PRDX1, with role is to reduce hydrogen peroxide, IFI30, a thiol reductase that can reduce protein disulfide bonds, hydroperoxides and GPX1 that catalyze the reduction of hydroperoxides in a glutathione-dependent manner to regulate cellular redox homeostasis(67) were all found more abundant in BPD. We imaged GPX1 in the alveolar parenchyma and found it present both in BPD and control in localizations consistent with Cap2 aerocytes. These cells are part of the air exchange interface and, as such, have likely developed molecular strategies to cope with the oxidative stress that occurs especially with premature delivery from the hypoxic fetal environment to relatively hyperoxic ambient and therapeutic supplemental oxygen and hyperoxemia. Based on the results of this study, the pulmonary endothelium appears exquisitely responsive to the redox environment as expected of cells at the air/blood interface responsible for gas transfer in the functional alveolar unit. A consequence of premature exposure to oxygen is dysregulation of capillary bed angiogenesis, as suggested by others(68). Interestingly, the donors in this study were 13-24 months old relatively remote from their neonatal hospitalization. Although some of the effects could be related to pre-mortem events, the consistency and apparent persistence of abnormal redox protein expression in the endothelium suggests that early oxygen exposures durably influence the longer-term response to oxidative stress. Future studies will be needed to elucidate whether the changes in protein abundance observed are specific to stable Cap2 cells or reflect changes in stem-cell states.

This study has a few key limitations. First, it was performed on tissue representing primarily the distal lung tissue and is not expected to reflect airway cellular alterations that contribute to BPD pathophysiology. A major limitation of the study is the heterogeneity of the analyzed cell populations that, although parent cell enriched, still contain mixtures of different cell types and subtypes assayed in bulk. BPD is inherently a heterogenous disease both within and between subjects. To gain a more detailed understanding of the roles of each cell type, further sorting into more defined subpopulations and refinement to single cell analyses will be necessary. At this time, we were not able to conduct such studies as, to our knowledge, no antibodies yet serve to sort pure populations of human alveolar epithelial (AT1 or AT2) and endothelial capillary cell types (Cap1 and Cap2). Another limitation is the low availability of human BPD tissues, especially in the context of the heterogenicity of the disease, as well as the costly and slow throughput of histologic imaging. While we provide data from 5 diseased and 5 aged- matched controls, immunofluorescence quantification demonstrated wide variability and did not always reach significance though clearly affected regions were apparent to visual inspection. A larger cohort of donors as well as advancing technology allowing broader regional, yet single cell analyses may well identify further differences between lung of children and adults born prematurely compared to those born at term. Due to the even lower availability of early-stage BPD tissues, in this study we opted for a description of later stage chronic BPD. Further studies, potentially with non-human primate models of BPD, are needed to understand how epithelial and endothelial cells contribute to BPD establishment and progression.

In conclusion, our proteomic approach provides valuable insights into key aspects of epithelial and endothelial cells in human BPD lungs. Similar studies performed on immune and mesenchymal populations could provide additional information on the role of these cells populations in BPD. Future research should explore other omics approaches, such as spatial proteomics, lipidomics, and metabolomics, along with post-translational modification (e.g., phosphoproteomics), to uncover and unite molecular-level changes in these cells for discovery in pathophysiology. Such efforts will dramatically enhance our understanding of the cellular and molecular mechanisms underlying BPD pathogenesis and could pave the way for developing targeted therapies to address the biological processes driving the disease.

## Acknowledgements.

This work was supported by a LungMAP consortium Pilot grant awarded to M.G. Ushakumary. LungMAP was supported by National Heart, Lung, and Blood Institute (NHLBI) Molecular Atlas of Lung Development Program Human Tissue Core (LungMAP HTC) grants U01HL122700 and U01HL148861 (to G.S. Pryhuber), and U01HL148860 (to J.P. Carson, J.N. Adkins, and G. Clair). Donor cell frozen vials were obtained from LungMAP BioRepository for INvestigation of Diseases of the Lung (BRINDL). We are grateful for the generosity of the donor families and honor their loss. Part of this work was performed in the Environmental Molecular Science Laboratory, a U.S. Department of Energy (DOE) national scientific user facility at Pacific Northwest National Laboratory (PNNL). Battelle operates PNNL for the DOE under contract DE- AC05-76RLO01830. The opinions expressed in this article are the authors’ own and do not reflect the view of the NIH, the Department of Health and Human Services, or the U.S. government.

---

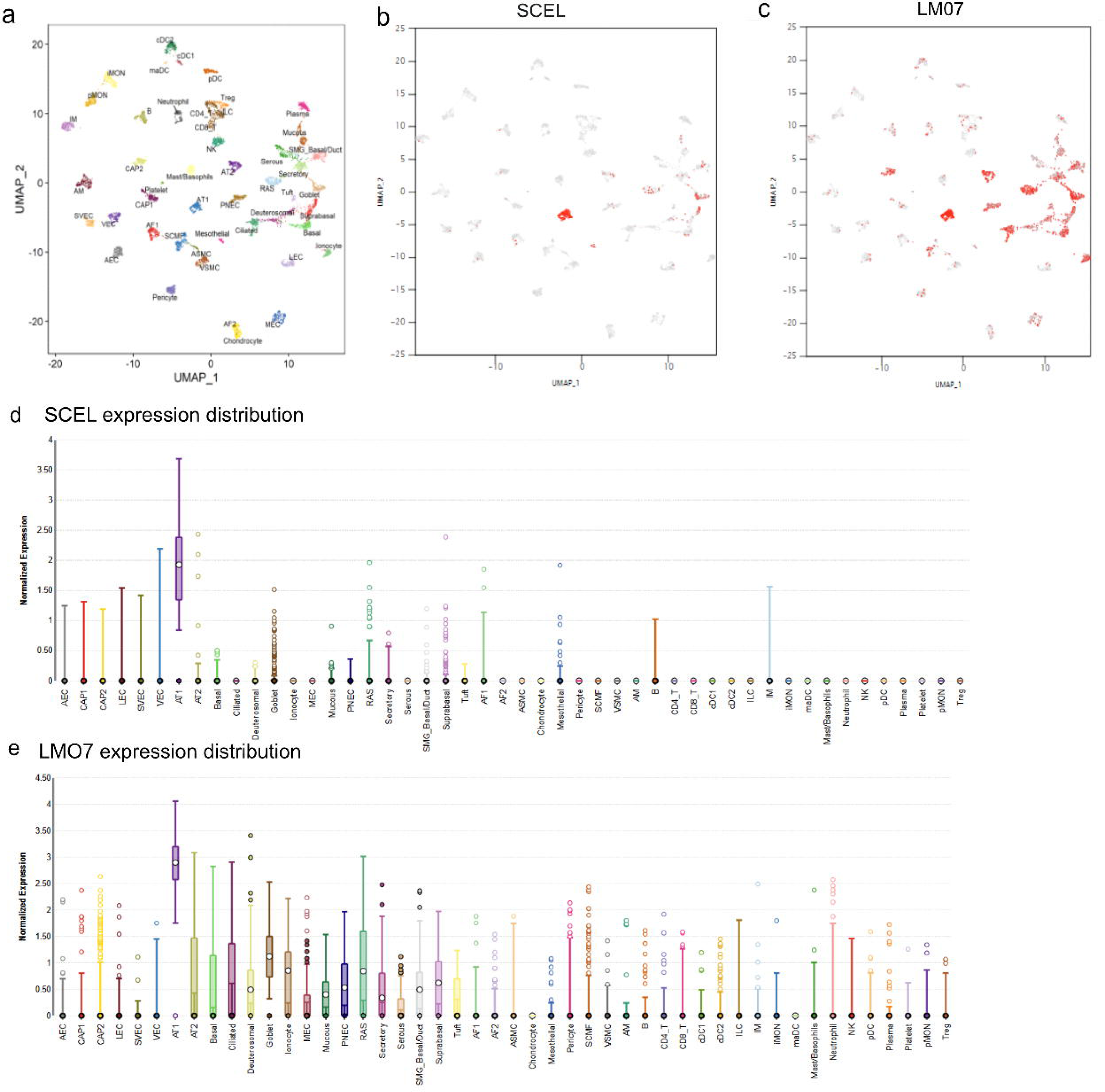

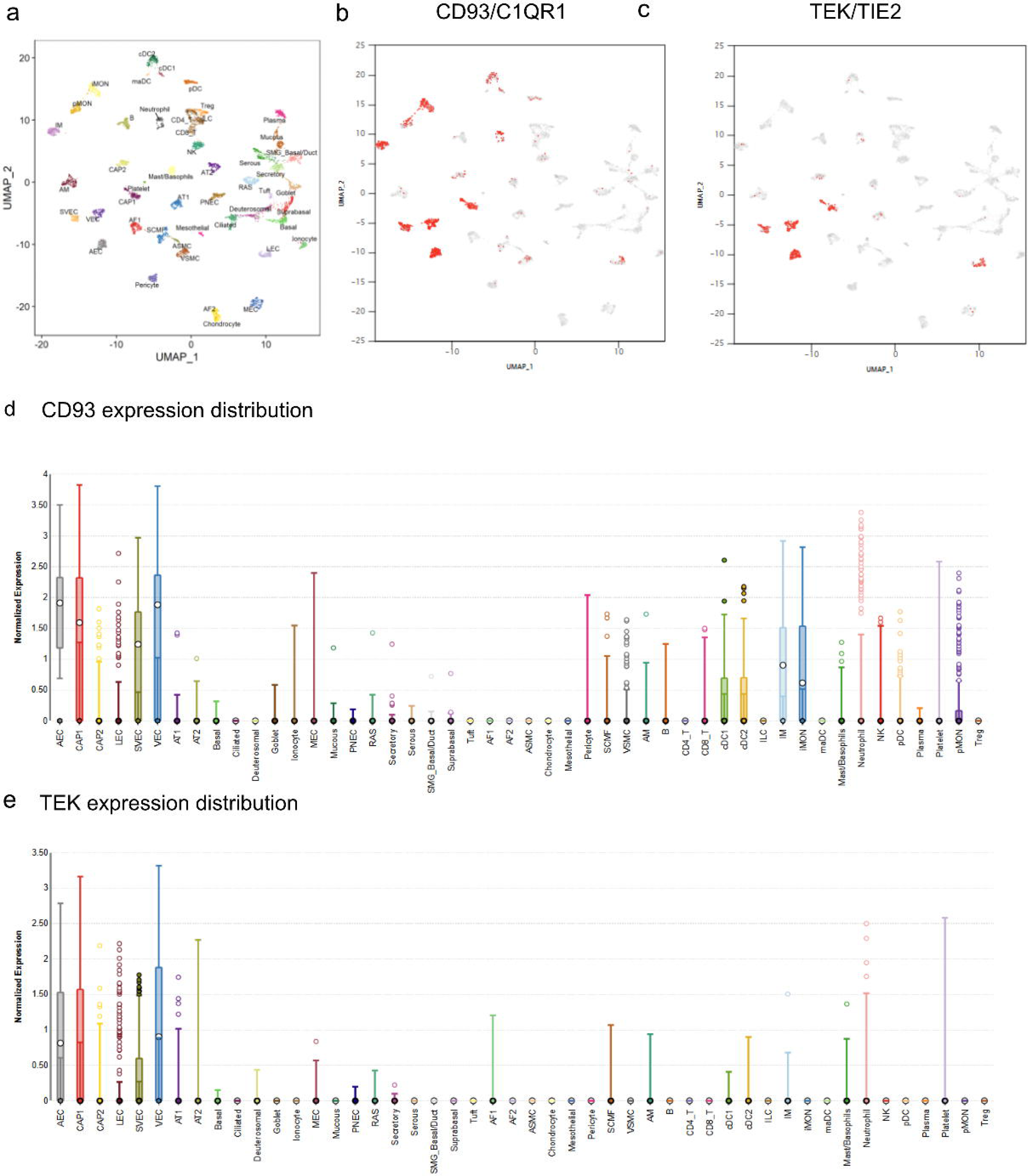

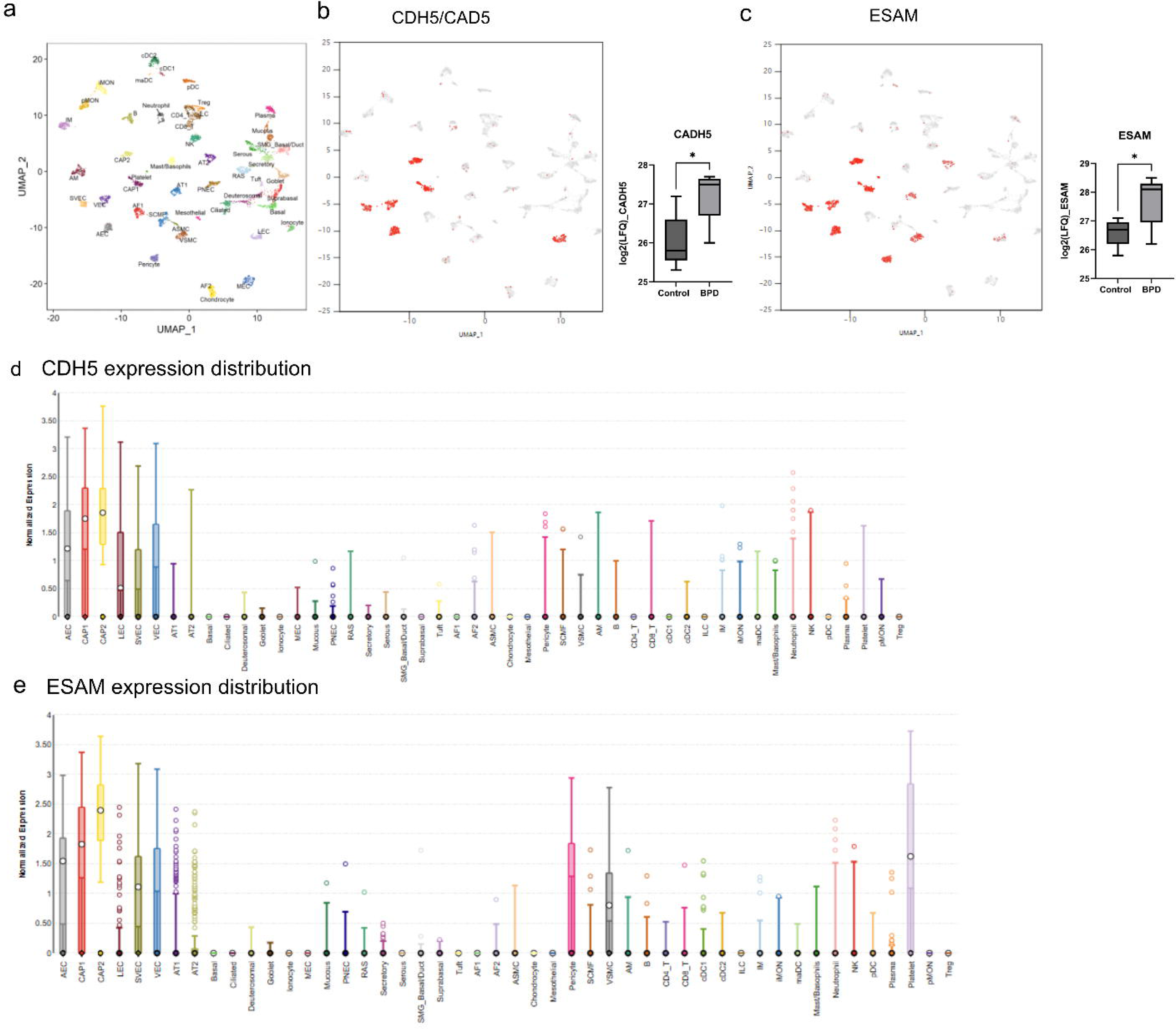

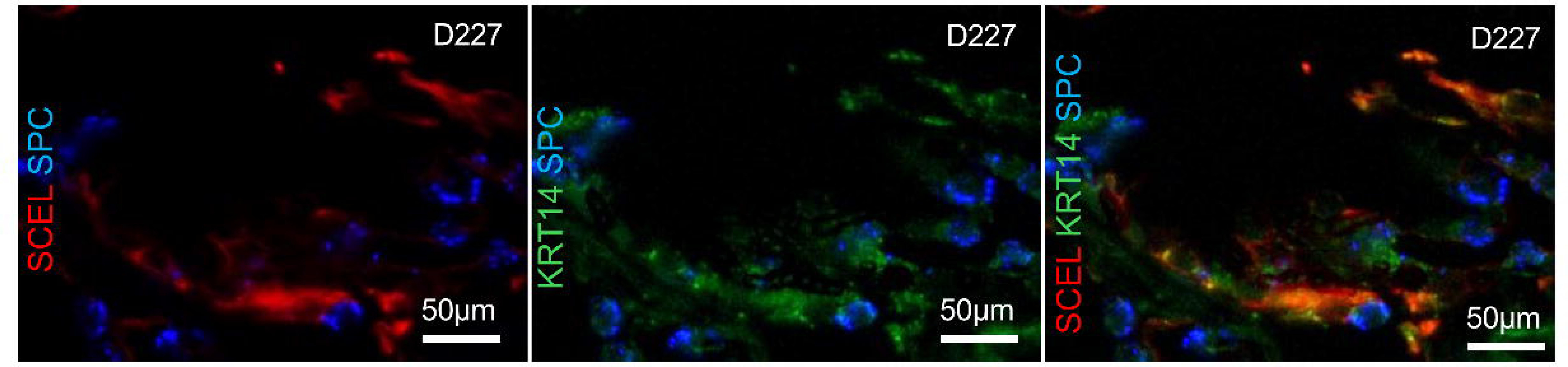

